# MOSAIC: a chemical-genetic interaction data repository and web resource for exploring chemical modes of action

**DOI:** 10.1101/112854

**Authors:** Justin Nelson, Scott W. Simpkins, Hamid Safizadeh, Sheena C. Li, Jeff S. Piotrowski, Hiroyuki Hirano, Yoko Yashiroda, Hiroyuki Osada, Minoru Yoshida, Charles Boone, Chad L. Myers

## Abstract

**Summary:** Chemical-genomic approaches that map interactions between small molecules and genetic perturbations offer a promising strategy for functional annotation of uncharacterized bioactive compounds. We recently developed a new high-throughput platform for mapping chemical-genetic (CG) interactions in yeast that can be scaled to screen large compound collections, and we applied this system to generate CG interaction profiles for more than 13,000 compounds. When integrated with the existing global yeast genetic interaction network, CG interaction profiles can enable mode-of-action prediction for previously uncharacterized compounds as well as discover unexpected secondary effects for known drugs. To facilitate future analysis of these valuable data, we developed a public database and web interface named MOSAIC. The website provides a convenient interface for querying compounds, bioprocesses (GO terms), and genes for CG information including direct CG interactions, bioprocesses, and gene-level target predictions. MOSAIC also provides access to chemical structure information of screened molecules, chemical-genomic profiles, and the ability to search for compounds sharing structural and functional similarity. This resource will be of interest to chemical biologists for discovering new small molecule probes with specific modes-of-action as well as computational biologists interested in analyzing CG interaction networks.

**Availability:** MOSAIC is available at http://mosaic.cs.umn.edu.

**Contact:**, or

## 1 Introduction

Discovering the mode-of-action of compounds is an important step in developing effective treatments for diseases or controlling the spread of pathogens. With the advent of efficient whole-genome sequencing methods that enable comprehensive genotyping of individuals, there are an increasing number of candidate drug targets implicated in specific diseases, which presents a need for new drug discovery paradigms that maximize these opportunities. The standard methods in high-throughput screening are relatively inefficient at linking small molecules to cellular targets and thus need to be improved in order to develop a rich collection of tool compounds for probing biological pathways and generating drug leads. We have yet to build a collection of small molecules that are able to target a substantial fraction of the human proteome.

Chemical-genomic approaches reveal compound-dependent phenotypes in a systematic, unbiased manner to predict the cellular role of an uncharacterized bioactive compound. For example, studies have tested compounds against the complete gene deletion collection that is available for the genetically tractable yeast *Saccharomyces cerevisiae* (Giaever et al., 1999; Parsons et al., 2004; Parsons et al., 2006; Hillenmeyer et al., 2008; Costanzo et al., 2010; Hoepfner et al., 2014; Lee et al., 2014; Giaever and Nislow, 2014; Wildenhain et al., 2016). In these approaches, a collection of genetic mutants is assayed for sensitivity or resistance to a compound to generate a compound-specific functional profile. We recently developed a new high-throughput platform for mapping CG interactions in yeast for large-scale compound screening and, using this, we generated a library of CG profiles for various compound collections (Piotrowski *et al.*, 2017). This represents one of the most extensive chemical-genomic datasets to date, including informative profiles for more than 1,000 natural products and derivatives, hundreds of clinically relevant compounds, and other small molecules. Here, we present MOSAIC, a web interface that provides access to this rich chemical-genomic dataset.

## 2 Methods

MOSAIC, the website for accessing the database, is located at http://mosaic.cs.umn.edu. This website displays data obtained by screening the RIKEN Natural Product Depository (NPDepo) as well as diversity libraries from the NCI/NIH, and the Published Kinase Inhibitor Set from Glaxo-Smith-Kline (Piotrowski *et al*., 2017). The data housed in the database were derived from the experiments described in (Piotrowski *et al.*, 2017) and were processed and analyzed using BEAN-counter (https://github.com/csbio/BEAN-counter) and CG-TARGET (Simpkins *et al.*, 2017) tools.

MOSAIC was designed using a combination of HTML, javascript with jquery, AJAX, javascript object notation (JSON), PHP, and MySQL. This provides a responsive environment for querying the CG interactions.

MOSAIC provides an interface for querying CG profiles, gene-and bioprocess-level target predictions. The MOSAIC interface has a search bar available in the middle or top of the browser that will accept a compound name, a gene common name or standard name, or a Gene Ontology (GO) biological process term. Multiple terms can be input by separating each term with a semicolon. The results page will return different data and predictions depending on whether a compound, gene, or process has been queried. If a query includes more than one type of input, the terms can be selected by using the tabs (Conditions, Genes, or GO Terms) and tables.

MOSAIC displays four relevant sets of information related to the mode-of-action of compounds. The first is the set of direct CG interactions, which reflect the sensitivity or resistance of each specific deletion mutant to a given compound. The second is the set of gene-level target predictions, which measure the correlation between the CG interaction profile and a genetic interaction profile. A high correlation indicates that the compound phenocopies the effect of a gene deletion, suggesting the compound perturbs the function of the corresponding gene. The third set of information is the bioprocess-level target prediction, which lists GO terms that are over-represented among the top gene-level target predictions as described in (Simpkins *et al.*, 2017). Finally, MOSAIC provides access to similarities between pairs of compounds based on either their chemical structure (Safizadeh *et al.*, 2017) or their CG interaction profile.

## 3 Conclusion

The MOSAIC database provides convenient access to CG interaction data, mode-of-action predictions, and chemical structural information for a large collection of both characterized and uncharacterized compounds. This resource can serve as a starting point for discovering new probes with specific modes-of-action or further characterization of novel compounds.

